# Impacts of the COVID-19 Pandemic on a Human Research Islet Program

**DOI:** 10.1101/2021.12.24.474114

**Authors:** Tina J. Dafoe, Theodore dos Santos, Aliya F. Spigelman, James Lyon, Nancy Smith, Austin Bautista, Patrick E. MacDonald, Jocelyn E. Manning Fox

## Abstract

Designated a pandemic in March 2020, the spread of severe acute respiratory syndrome virus 2 (SARS-CoV2), the virus responsible for coronavirus disease 2019 (COVID-19), led to new guidelines and restrictions being implemented for individuals, businesses, and societies in efforts to limit the impacts of COVID-19 on personal health and healthcare systems. Here we report the impacts of the COVID-19 pandemic on pancreas processing and islet isolation/distribution outcomes at the Alberta Diabetes Institute IsletCore, a facility specialising in the processing and distribution of human pancreatic islets for research. While the number of organs processed was significantly reduced, organ quality and the function of cellular outputs were minimally impacted during the pandemic when compared to an equivalent period immediately prior. Despite the maintained quality of isolated islets, recipient groups reported poorer feedback regarding the samples. Our findings suggest this is likely due to disrupted distribution which led to increased transit times to recipient labs, particularly those overseas. Thus, to improve overall outcomes in a climate of limited research islet supply, prioritization of tissue recipients based on likely tissue transit times may be needed.

## Introduction

Human pancreatic tissue and isolated islets of Langerhans are vital to research, where they are used to study islet morphology, β-cell proliferation, genomics, insulin and glucagon secretion, fuel-induced toxicity, transcription factor regulation, transplantation, and many other aspects of endocrine physiology and diabetes. Characterization of human islet function is central to understanding diabetes, given the key role that islets play in disease pathophysiology^1^ and genetic susceptibility^2–4^, and to regenerative strategies including the production of mature human β-cells from stem cells^5–7^. Increased interest in—and need for—these areas of research, along with awareness of the limitations of non-human models, has elevated the demand for human research islets. Concerns have been raised regarding future access^8,9^ with data from the European Consortium for Islet Transplantation (ECIT) suggesting a growing gap between human research islet supply and demand^10^, a trend first reported by the Islet Cell Resource Consortium (the predecessor to the Integrated Islet Distribution Program, or IIDP).^11^

To address the growing demand for human research tissue, the Alberta Diabetes Institute (ADI) IsletCore was established in 2010 with the goal of isolating, distributing, and biobanking insulin-producing pancreatic islets from donor organs with research consent, but not accepted into clinical transplantation programs^12^. The ADI IsletCore is located at the University of Alberta in Edmonton and is one of the world’s largest programs of islet isolation and distribution exclusively for research, providing services to over 130 research groups globally. Over the past 11 years, this program has grown to include the provision of biobanked samples, including FFPE sections; cryopreserved and snap frozen islets; and custom collection of additional pancreas-associated tissues including spleen, adipose, intestine, lymph nodes, and blood^13^; with public sharing of data on basic donor information, quality measures, biobank inventory, and functional analysis (**Figure 1**)^14^.

**Figure 1.**
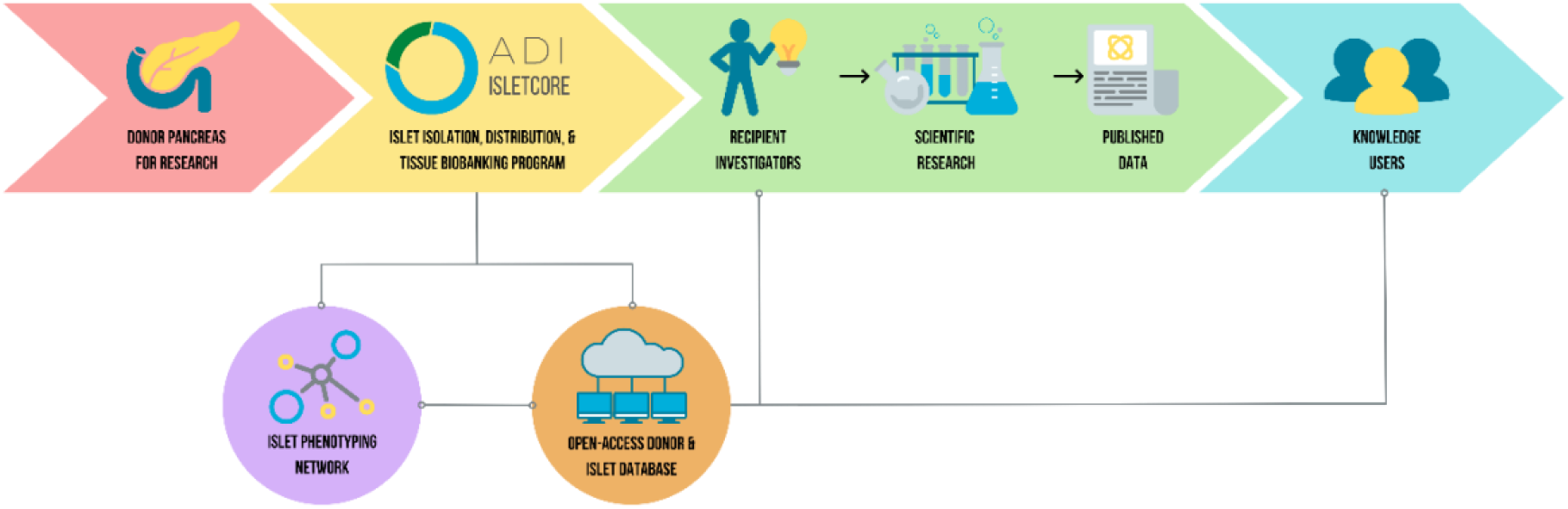
Schematic illustration of ADI IsletCore operational flow.

The World Health Organization (WHO) declared COVID-19 a pandemic on March 11^th^, 2020^15^ and ADI IsletCore opted to halt processing of pancreases at this time. On March 17^th^, 2020 the Government of Alberta declared a Provincial State of Public Health Emergency under the *Public Health Act* for the first time in its history^16^, and the University of Alberta implemented non-essential research restrictions, with buildings closed to all but essential and COVID-19-related research on March 23^rd^, 2020. Subsequent additional measures included travel restrictions and quarantine requirements, limitations on gatherings, mandatory two-metre physical distancing, and mask wearing requirements in indoor settings^16^. Similar restrictions were implemented around the world, as jurisdictions strived to protect their communities and healthcare systems^17^.

In line with a nationwide implementation of duplicate negative SARS-CoV2 testing for organ donation, and following implementation of revised standard operating procedures that enabled our islet isolation team to work safely, the ADI IsletCore was able to resume operations on May 25th. Unfortunately, this two-month closure of the ADI IsletCore was the first of several, with three subsequent temporary shutdowns coinciding with ongoing waves of COVD-19 cases, hospitalisations, and intensive care unit admissions experienced within Alberta^18^ (**Figure 2a**). While the provincial- and university-mandated research restrictions did not impose the closure of the ADI IsletCore, this action was taken to limit impact on the healthcare system, which was under significant strain during these times. Coordinating organ procurement on our behalf impacts the workload of Organ Procurement Organization staff, as well as necessitating multiple contacts between healthcare staff and transportation service providers. We chose to halt our acceptance and processing of pancreases, while maintaining user access to our biobanked samples. In this report, we assess the overall impacts of the COVID-19 pandemic on the operations and outcomes of the ADI IsletCore facility.

**Figure 2.**
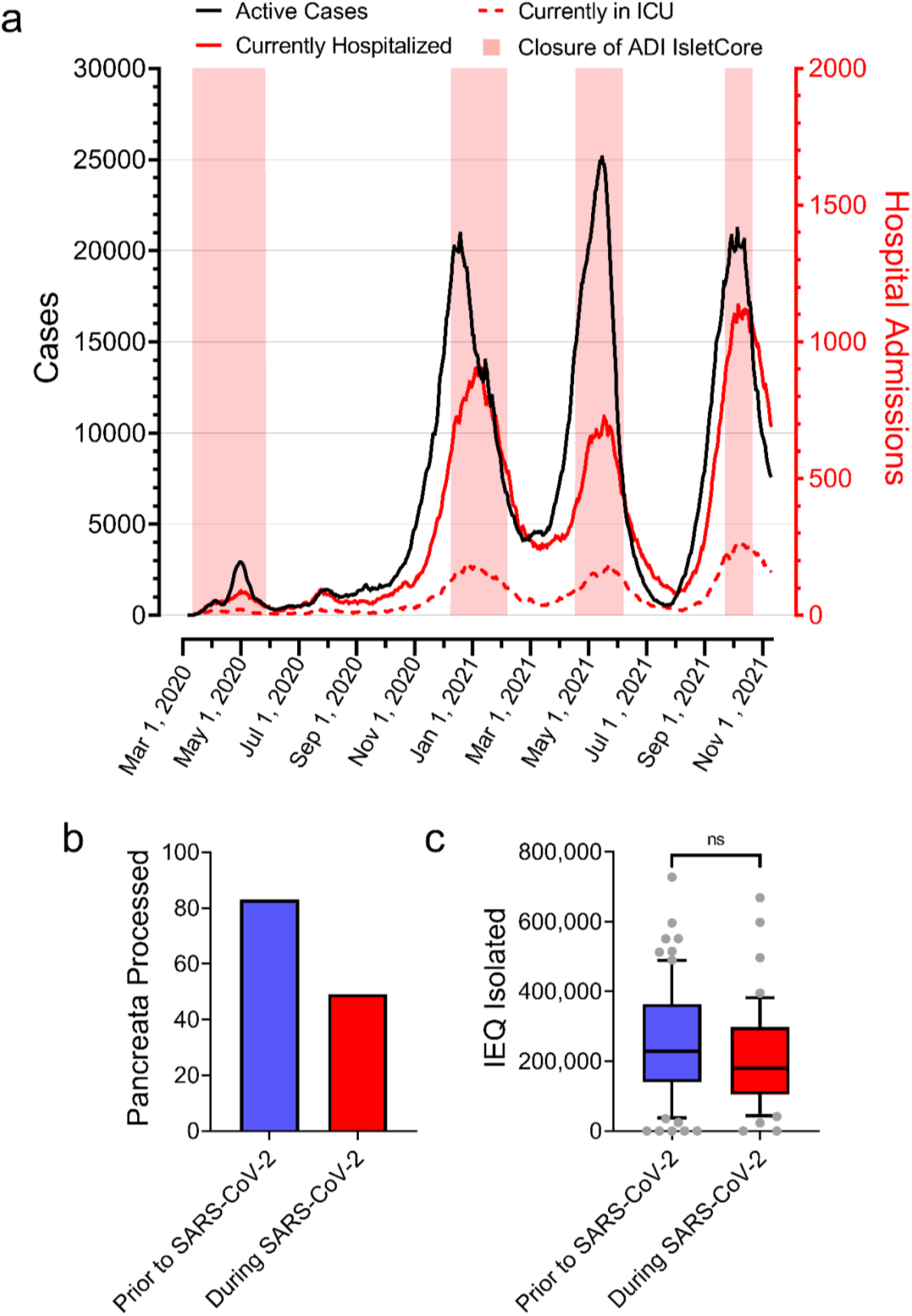
(a.) Active COVID-19 case counts, hospitalizations, and Intensive Care Unit (ICU) admissions in Alberta and ensuing operational closure of ADI IsletCore (pink). (b.) Pancreata processed and (c.) number of islet equivalents (IEQ) isolated prior to, and during the COVID-19 pandemic.

## Methods

### Data sets and statistics

To report the impacts of the COVID-19 pandemic on ADI IsletCore’s operations, we reviewed data held in our database (REDCap 11.1.24) comparing two 19-month time periods: one beginning on the date of the ADI IsletCore’s first closure (March 12^th^, 2020, to October 12^th^, 2021: during COVID-19, abbreviated hereafter as DC), and one immediately prior to (September 12^th^, 2018, to March 11^th^, 2020; pre-COVID-19, abbreviated hereafter as PC). Comparison of continuous parameters between the two timeframes was assessed by unpaired t-test. Non-continuous parameters were compared via a Chi-square test. A probability value of p<.05 was considered statistically significant.

### Pancreas Procurement

The ADI IsletCore processes pancreases from human organ donors across Canada. Pancreases are offered to ADI IsletCore only when they are deemed not ideal for clinical transplant purposes and proceeds only in cases of retrieval of other abdominal organs for transplantation. Consent for research use of the pancreas and associated tissues is obtained by the local Organ Procurement Organizations, who coordinate the donation process. All donors undergo serology testing for infectious agents, including SARS-Cov2, prior to donation. We process pancreases that have been donated following neurological determination of death (NDD), circulatory death (DCD), or medical assistance in dying (MAID). Organs are expedited in University of Wisconsin organ preservation solution (UW) or equivalent (HTK, SPS) on ice to our facility using courier service and a commercial airline (Air Canada). In situations where other organs are being transported to Edmonton, pancreas may be transported with these by a medical flight. Locally procured pancreases are collected directly from the University of Alberta Hospital.

### Islet Isolation

The islet isolation process is adapted from that of Ricordi *et al*^19^ and involves both mechanical and enzymatic digestion of the pancreas. The organ is perfused with collagenase and neutral protease enzymes via the pancreatic ductal system, the tissue is agitated using a Ricordi chamber and auto-isolator, and the released islets are separated via a stainless-steel mesh. Subsequent purification of the digested tissue is performed via continuous gradient centrifugation and fraction collection. Full protocol details are publicly available via protocols.io^20^.

### Islet Quantification and Purity Assessment

Human islets are quantified by using a microscope with an eyepiece reticle, in conjunction with a grid placed under the sample to ensure that each islet is only counted once. The islet sample is first stained with the zinc-binding dye, dithizone (DTZ), which enables visual identification of islets by their red colour. Acinar tissue, which lacks zinc-containing beta cells, remains unstained. Preparation purity is estimated by visualizing the percentage of the sample that is DTZ-positive. Volumetric adjustment of human islets of different sizes to standardized Islet Equivalents (IEQ) is performed, and all islet quantifications are reported in IEQ^**20**^.

### Insulin and DNA content

Islet preparations are characterized with respect to total cellular insulin (Alpco, Salem, NH, USA), and DNA content (Quant-iT™ PicoGreen ^®^ dsDNA, Molecular Probes, Eugene, OR, USA) following the manufacturers’ instructions^20^.

### Glucose-Stimulated Insulin secretion

Insulin secretion measurements are performed at 37°C in Krebs Ringer Buffer (KRB) (in mM: NaCl 115; KCl 5; NaHCO_3_ 24; CaCl_2_ 2.5; MgCl_2_ 1; HEPES 10; 0.1% BSA, pH7.4) with glucose concentrations as noted. Triplicate groups of fifteen islets are pre-incubated for two hours with 1 mM glucose KRB. Islets are subsequently incubated for one hour in 1 mM glucose KRB followed by a one-hour stimulation in either 10mM or 16.7 mM glucose KRB. Supernatants are removed and total insulin content extracted from the islet pellet using acid-ethanol. Samples are stored at −20°C and assayed for insulin via chemiluminescence (Alpco, Salem, NH, USA)^20^.

### Islet Distribution

Prior to distribution, islets are cultured in supplemented Connaught Medical Research Laboratories (CMRL) media at 22°C, 5% CO_2_ for approximately 12 to 96 hours, depending on time of the isolation. On the day of distribution, IEQs are re-quantified and samples are aliquoted into 50 ml tubes, brought to volume with CMRL media, and packaged for shipment at ambient temperature. Tubes are packaged on their side to prevent islet pelleting, and a temperature indicator is included with the shipment. Islets are distributed via FedEx overnight service within Canada and the USA, with shipments overseas typically delivered within 48 hours.

### Recipient Feedback

Feedback on each islet preparation received is requested from all recipient groups using an online form. Quality is assessed via multiple choice options of excellent, good, fair, or poor. User location is noted, and specific comments or concerns can be reported.

## Results

### Organ processing

The ADI IsletCore’s islet isolation activities were on hold for a total of 34 weeks, or 41% of the 19-month COVID-19 period during which we were unable to process organs (**Figure 2a**). Accordingly, a reduction in the number of organs processed during the pandemic was observed, dropping from 83 pre-COVID-19 (PC) to 49 during COVID-19 (DC) (**Figure 2b**), a similar 41% reduction in organs processed.

As expected, the reduced number of organs received was paralleled by a decreased total yield of islets isolated (**Figure 2c**; 19.4 million IEQ PC versus 9.9 million IEQ DC). However, islet isolation success is also dependent on the quality of the donor pancreas^21^, with factors such as age, BMI, medical history, and type of donation all having an impact on islet yield and function^12,22–28^. We compared each of these criteria between the two timeframes of interest, to determine whether these were altered DC (**Figure 3**). Donor age was significantly lower (47.6 +/− 1.7 years PC versus 41.7 +/− 2.4 years DC, p<.05), whereas donor sex, BMI, HbA1c, and type of donation (NDD, DCD or MAID), pancreas mass and circulating lipase and amylase did not differ between the two timeframes. Although the percentage of donors for whom over-dose or motor vehicle accident was recorded as the cause of death is reported to have increased and decreased, respectively, during the pandemic^29^, we find no significant changes in the causes of death for donors included in our study (data not shown). Ultimately, the mean number of IEQ isolated from each donor (252247.8 IEQ PC +/− 17771.6 IEQ PC versus 214179.6 +/− 21285.7 IEQ DC, n=49-83, p=NS) and per gram of pancreas weight (2945.4 +/− 226.8 IEQ/g PC versus 2333.4 +/− 211.2 IEQ/g DC, n=49-83, p=NS) remained unchanged, indicating that, despite the observed decrease in donor age, the reduction in total IEQ yield during the COVID-19 period appears to be entirely accounted for by a reduction in pancreases processed.

**Figure 3.**
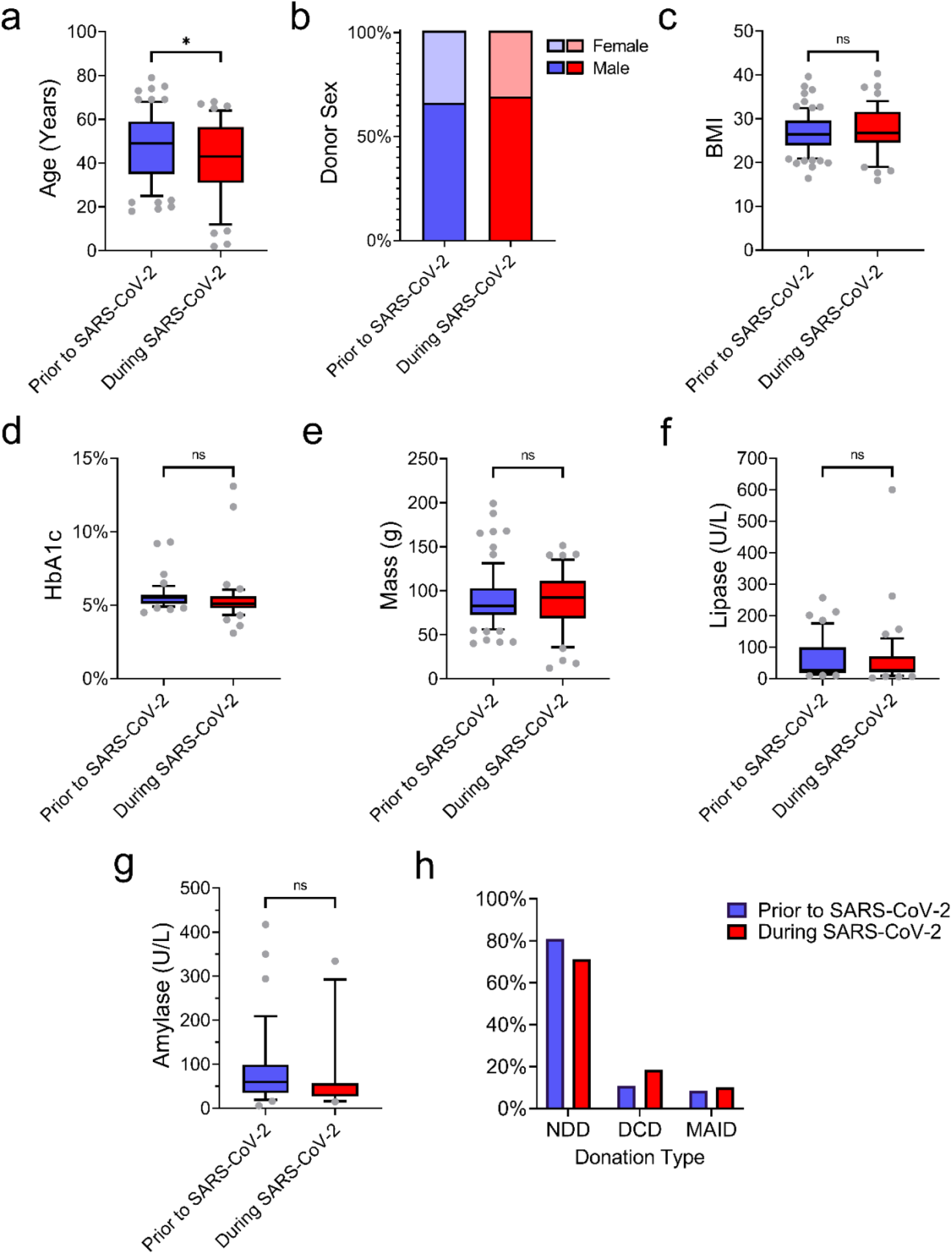
Donor parameters prior to (blue) and during (red) SARS-CoV2. (a.) Donor age. (b.) Donor sex. (c.) Donor body mas index (BMI). (d.) Donor glycated hemoglobin (HbA1c). (e.) Pancreas mass. (f.) Donor circulating lipase level. (g.) Donor circulating amylase level. (h.) Type of donation. N=11-83. *p<0.05.

### Logistics

The time between pancreatic cross-clamp in the operating theatre and the start of islet isolation at the ADI IsletCore, governed by surgical, transportation and staffing schedules, is referred to as cold ischemia time (CIT). Extended CIT is known to have an adverse influence on islet isolation^12^. As such, ADI IsletCore typically only accepts donors with a cold ischemia time of less than 24 hours and historically has a program-wide mean CIT duration of 14.4 +/− 0.3 hrs.

Global supply chain and transportation logistics issues have been a notable result of the pandemic, impacting many sectors. ADI IsletCore receives pancreases from across Canada, which are transported to Edmonton via a single airline. A Government of Canada travel advisory to avoid non-essential travel^30^, combined with measures to restrict the transmission of COVID-19 on domestic and international flights^31^, led to a reduction in scheduled commercial flight activity across the country^32^. We were surprised to find that there was no resultant increase in the CIT (**Figure 4a**), and instead a significant decrease was observed (14.4 +/− 0.6h PC versus 11.6 +/− 0.8h DC, n=46-83, p<.005). While transit times from most provinces remained unchanged, an overall decrease in CIT was primarily driven by a reduction in CIT for local Alberta donors (11.6 +/− 1.1 hrs PC versus 6.6 +/− 1.1 DC, n=18-19, p<0.005). This is most likely due to a sense of urgency to process pancreases as soon as possible, regardless of antisocial work hours, due a perceived increase in value of each donor at this time. An observed increase in the percentage of donors obtained locally (24.1% PC vs 36.7% DC) and decrease in those obtained from centres east of Ontario (22.9% PC versus 12.2% DC) is likely reflective of the logistical impacts of obtaining organs from out-of-province and contributes to the reduced overall CIT (**Figure 4b**).

**Figure 4.**
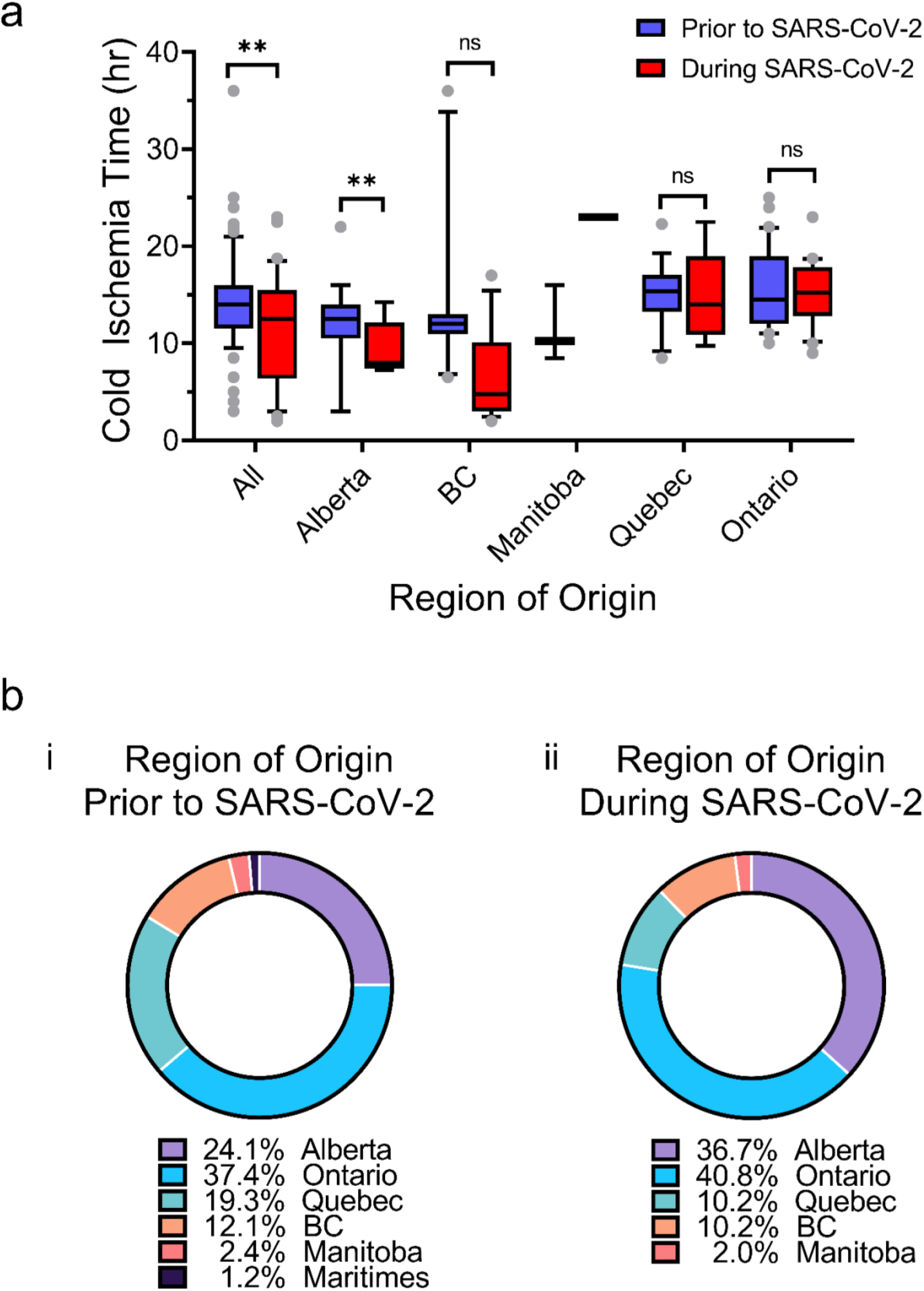
(a.) Cold ischemia time for pancreases transported from each of the organ procurement regions prior to (blue) and during (red) SARS-CoV2 (**p<0.005. n=2-31 for each region). (b.) Breakdown of regions from which pancreases were received prior to (i) and during (ii) SARS-CoV2.

### Islet Distribution

The ADI IsletCore’s primary goal is to provide human islets to the broader research community. Currently we serve 132 research groups, 50% of which are in the US, 25% in Canada, and 25% in Europe, Asia or Australasia. Despite the observed reduction in the number of pancreases processed and islets isolated, we were able to fulfil 86% of requests for islets, both prior to and during COVID-19. ADI IsletCore uses FedEx courier service to distribute samples to recipient labs. Courier services were affected by government restrictions to contain the spread of COVID-19, and experienced increased volumes, as well as disruptions caused by COVID-19 outbreaks and severe weather events at hub locations. These factors combined to result in additional transit time and frequent delivery delays and continue to impact human islet shipments at the current time (November 2021). While 94.5% of shipments within Canada met the standard of next day delivery, 13.5% of those to the United States were delayed, and 43.9% of international shipments failed to arrive with the standard of 48 hours (**Figure 5b**). In several cases, international transit times were greater than five days, with several shipments taking seven or eight days to reach their destination.

**Figure 5.**
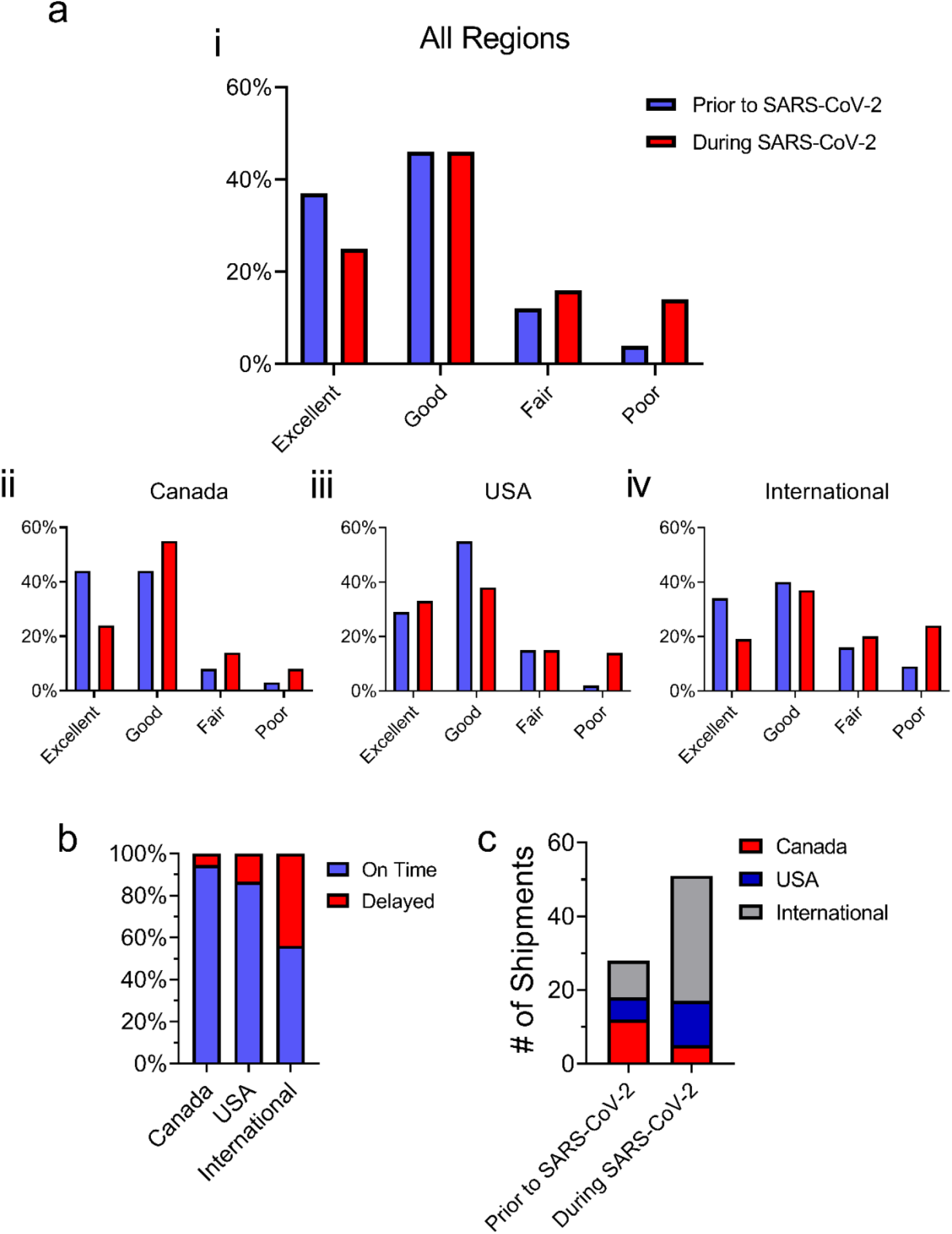
Feedback obtained from recipient groups regarding the perceived quality of islets. (a.) Total feedback ratings are shown for the period prior to (blue) and during (red) SARS-CoV2 (i), as well as those received from recipient groups located in Canada (ii, n=131-201), USA (iii, n=79-128) or internationally (iv, n=70-99). (b.) Percentage of shipments to different geographic areas that were delivered within the standard delivery times (n=107-162). On-time shipments are shown in blue, delayed deliveries are indicated in red. (c.) Number of shipments for which refunds were requested due to perceived low islet quality from each geographic area during the period prior to and during SARS-CoV2. Canada is represented in red, USA in blue, and international groups are shown in grey.

Feedback from recipient labs is an important tool in improving the quality of islets and efficiency of shipping procedures provided by ADI IsletCore and is publicly available (www.isletcore.ca). We observed an overall decrease in the reported quality of our preparations during COVID-19, compared to prior to the pandemic (**Figure 5ai**). In all geographic areas, a decrease in the percentage of preps rated “excellent” or “good” was observed, and an increase in “fair” and “poor” preps was reported (**Figure 5aii, iii, iv**). This was also reflected in an increase in the number of requests for refunds (9% IEQ refunded DC versus 2% IEQ refunded PC). There was also a notable increase in the refund requests relating to international shipments (**Figure 5c**). While these international shipments accounted for only 28.2% of the total, they represented 67.7% of the refund requests DC, an increase from 35.7% PC.

### Islet Function

Although increased shipping times likely accounts for much of the increase in negative feedback, we wondered whether poorer islet outcomes and performance may be driving some decrease in the perception of islet quality. Insulin secretion in response to 1 mM, 10mM and 16.7 mM glucose is measured for each prep as part of our standard quality assurance and human islet phenotyping program. The total islet insulin content (**Figure 6a**), and insulin secretory response to glucose, whether expressed as absolute values (**Figure 6bi**), percent content (**Figure 6bii**) or stimulation index (**Figure 6biii**), was unchanged during the pandemic. Other in-house parameters of islet quality and function were similarly unchanged (**Figure 7**) including preparation purity, percentage of islets trapped in acinar, islet particle index, and insulin content per IEQ. The mean duration in culture prior to distribution and the percentage recovery following culture were also unchanged. Together, these findings suggest that the perceived reduction in the quality of islets reported by recipient groups was likely due to shipping disruptions and the ensuing extended duration that islets were subjected to suboptimal culture conditions during transit^33–36^.

**Figure 6.**
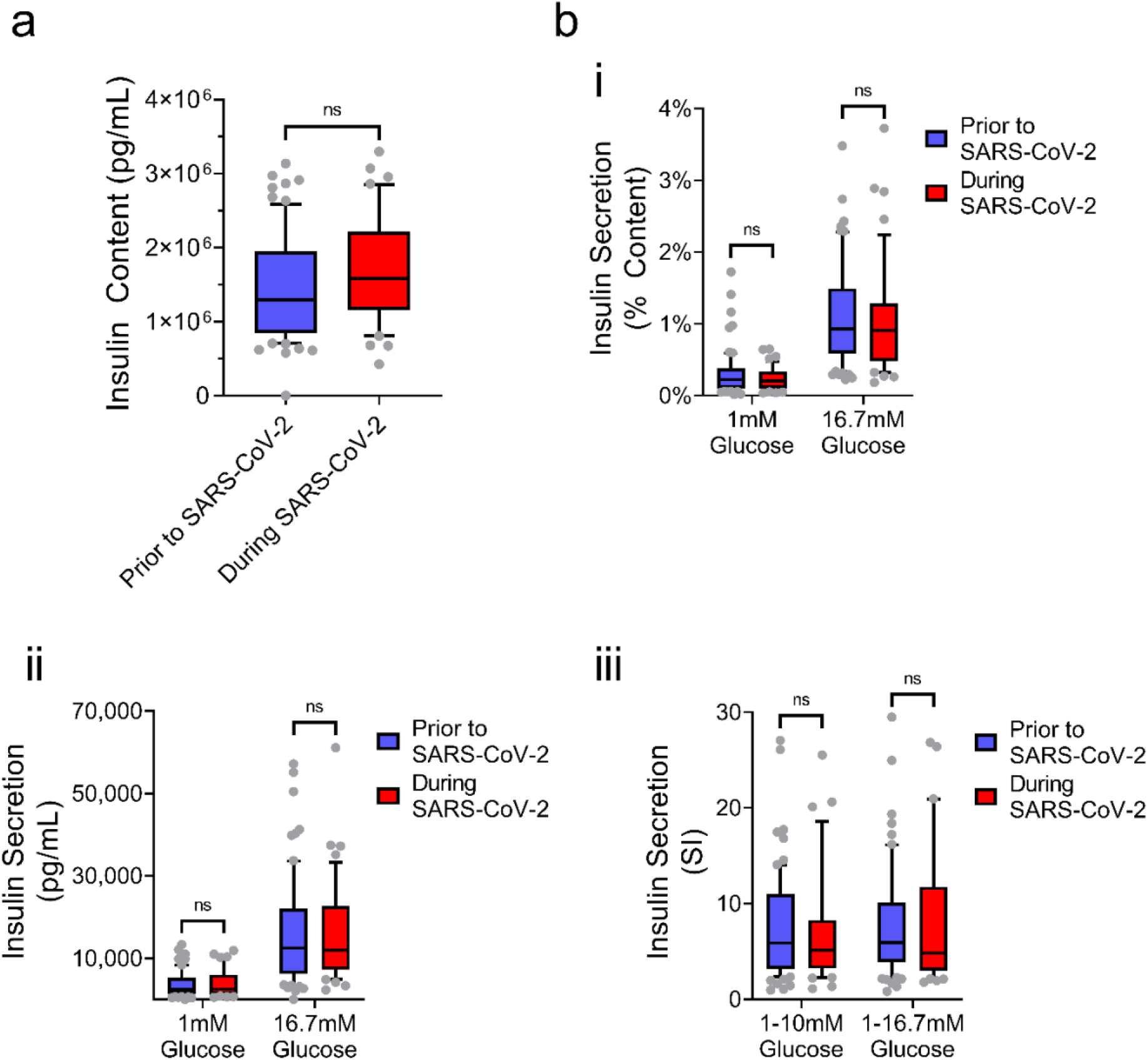
Measures of islet insulin secretory responses to a glucose challenge, from human islets isolated pre-COVID-19 (blue) and during COVID-19 (orange). (a.) Islet insulin content. (b.) Islet insulin secretion expressed absolute values (i), percent content (ii) and stimulation index (iii). Data represents 128-209 measurements from 43-70 donors per group.

**Figure 7.**
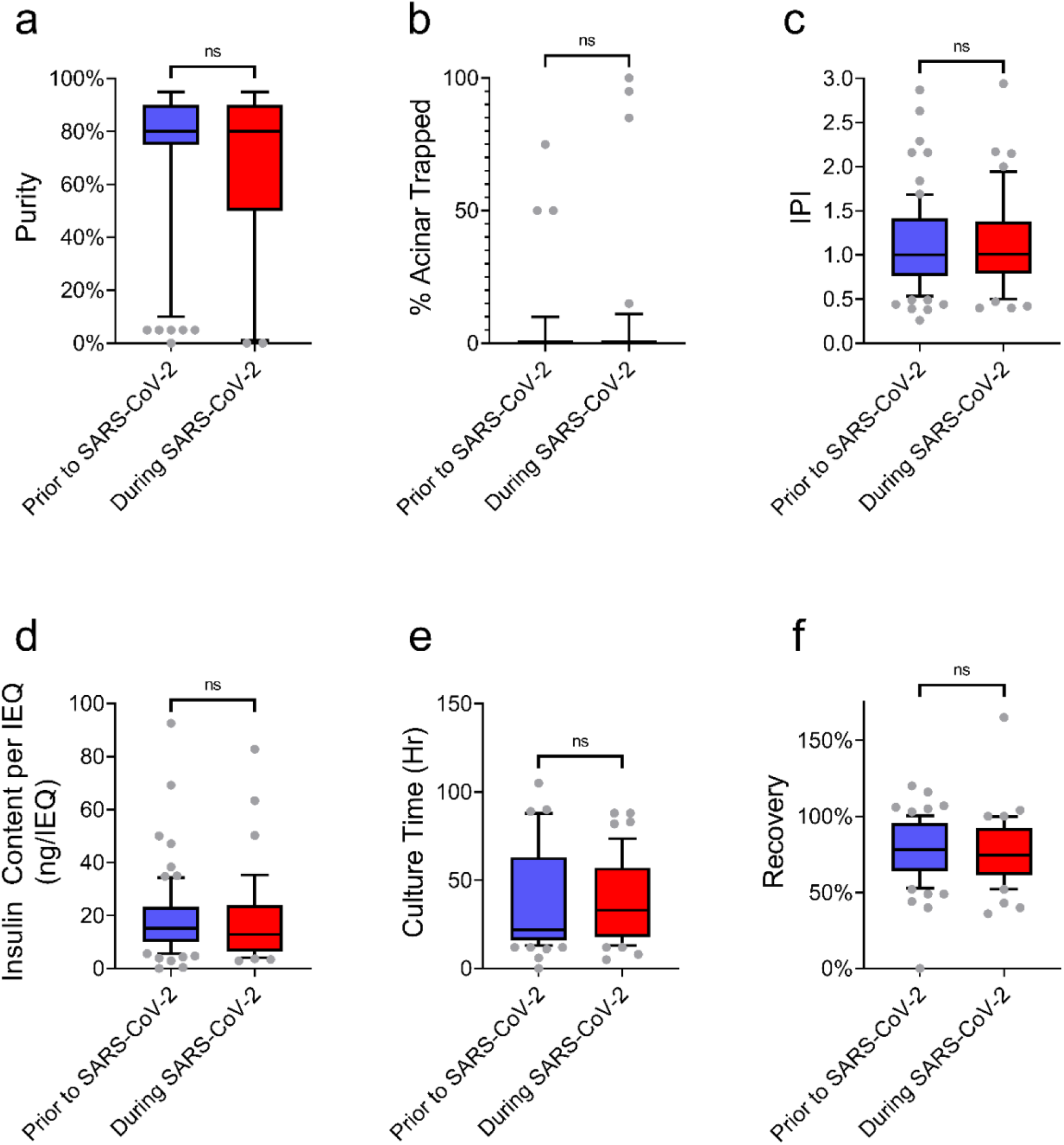
Islet isolation outcomes and quality control parameters prior to and during the COVID-19 pandemic. (a.) Islet purity. (b.) Percentage of islets remaining trapped in acinar tissue following isolation. (c.) Islet Particle Index (IPI). (d.) Insulin content per islet equivalent. (e.) Duration of islet culture, prior to distribution of islet to recipient groups. (f.) Percentage of islets recovered, following culture. n=49-83 islet isolations.

## Discussion

The importance of accessible and transparent collecting and reporting of human islet donor characteristics and phenotyping has recently been highlighted^8^ and is now a requirement for publication in major journals in the field^37,38^. Variables such as donor age, BMI, medical history, and type of donation have been previously demonstrated to impact islet yield and function^12,22–28^. As such, ADI IsletCore captures metadata encompassing 64 variables pertaining to each of our donors^14^, hosted locally within the University of Alberta Faculty of Medicine & Dentistry’s secure data centre. Technical parameters related to the isolation process, such as cold ischemia time^12,22,26^, can also influence islet yield and function^23,27,28^, and 56 datapoints describing these variables, plus quality assurance assessments and functional phenotyping of the islets, are also captured in our database. As such, the ADI IsletCore’s extensive database of donor, isolation, and functional parameters relating to its islet isolation activities presents an opportunity and resource for retrospective assessment of the impacts of the COVID-19 pandemic on our research islet isolation program.

The primary impact of the pandemic was closure of the ADI IsletCore on multiple occasions over the past 19 months, amounting to 41% of the pandemic period included in this study. The decision to suspend activities multiple times during the pandemic was challenging, given the impacts of this not only on our own staff and research, but for the many research groups worldwide that rely on human tissue for their own studies. ADI IsletCore staff did not remain idle during facility closures, and instead took the opportunity to review our operational and administrative procedures, including the production of the “ADI IsletCore Welcome Booklet”, a novel resource which serves as a “User Guide to Human Islets” for islet biologists^39^.

While the closures of our facility had major impacts on the number of organs processed and islets isolated, we found the reduction in organs processed to be less than expected. Given previous reports describing redirection of resources, increased screening requirements, decreased donations, and suspension of transplantation programs^29,40–42^, we expected a reduction in pancreases received during the periods when the ADI IsletCore was operational. However, our experience did not reflect the reported impacts on organ donation, and we continued to receive donor pancreases at the same frequency as during pre-pandemic times. Similarly, previously-reported impacts on donor cause of death^29^ were not reflected in our study. The organs offered to our program constitute only a subset of donated organs however, since organ priority is given to clinical transplantation programs and research programs that are geographically proximal to the donating centre. It is possible that the characteristics of the subgroup of organs in our study might not be indicative of the impacts of COVID-19 on the Canadian organ donation system at large.

The steady influx of pancreases during our operational periods enabled us to maintain our pre-COVID-19 level of request fulfillment. It is likely that any build-up of demand for islets following ADI IsletCore closure periods was counter-balanced by restrictions placed on recipient researcher centres, including facility closures, a necessity for split shifts, staff redeployments and staffing furloughs or absences^43^. The balanced level of demand, in proportion to the number of pancreases processed, also emphasizes the requirement for islets from multiple donors in research, rather than a total quantity of islets from a single or few donors. While in-house measures of islet quality and function remained unchanged, it is apparent that significant deterioration of islet quality occurred during transit, which is reflected in the reduced positive feedback from recipient groups. Unfortunately, extended transit times for islet shipments became a common occurrence during COVID-19, with FedEx suspending guaranteed delivery standards and releasing a service update stating, “due to ongoing impacts from the COVID-19 pandemic, we are experiencing significant volumes, which may result in potential transit delays in your area.”^44^

Fortunately, the COVID-19 pandemic has not prevented ongoing and exciting research in islet biology, with many papers describing important advances being published during this unusual period^45^. Following the period reviewed in this study the ADI IsletCore has resumed full operations, although potential impacts of future COVID-19 waves remain unknown. At the time of writing, 85.2% of Alberta’s population aged 12 years and over is fully vaccinated, a requirement for several activities and workplaces. We are optimistic that with continued caution, government regulation, and vaccination, that the ADI IsletCore will be able to operate effectively in service of the diabetes research community going forward.

## Acknowledgments

We thank the Human Organ Procurement and Exchange (HOPE) program and Trillium Gift of Life Network (TGLN) for their work in procuring human donor pancreas for research. We especially thank the organ donors and their families for their kind gift in support of diabetes research. Establishment of the ADI IsletCore was initially supported by funding from the Alberta Diabetes Foundation and the University of Alberta. Access to, and maintenance of, database resources is supported by the Women & Children’s Health Research Institute (WCHRI) in collaboration with the Northern Alberta Clinical Trials and Research Centre (NACTRC). TdS is supported by an International Helmholtz Research School for Diabetes Scholarship, CIHR Frederick Banting and Charles Best Canada Graduate Scholarship, the Walter H Johns Graduate Fellowship from the University of Alberta, and an Alberta Innovates – Data Enabled Innovation Scholarship. PEM holds the Canada Research Chair in Islet Biology.

## References

1. Kahn SE, Cooper ME, Del Prato S. Pathophysiology and treatment of type 2 diabetes: perspectives on the past, present, and future. Lancet Lond Engl 2014; 383:1068–83.

2. Dimas AS, Lagou V, Barker A, Knowles JW, Mägi R, Hivert M-F, Benazzo A, Rybin D, Jackson AU, Stringham HM, et al. Impact of type 2 diabetes susceptibility variants on quantitative glycemic traits reveals mechanistic heterogeneity. Diabetes 2014; 63:2158–71.

3. Ingelsson E, Langenberg C, Hivert M-F, Prokopenko I, Lyssenko V, Dupuis J, Mägi R, Sharp S, Jackson AU, Assimes TL, et al. Detailed physiologic characterization reveals diverse mechanisms for novel genetic Loci regulating glucose and insulin metabolism in humans. Diabetes 2010; 59:1266–75.

4. Olsson AH, Volkov P, Bacos K, Dayeh T, Hall E, Nilsson EA, Ladenvall C, Rönn T, Ling C. Genome-wide associations between genetic and epigenetic variation influence mRNA expression and insulin secretion in human pancreatic islets. PLoS Genet 2014; 10:e1004735.

5. Bruin JE, Saber N, Braun N, Fox JK, Mojibian M, Asadi A, Drohan C, O’Dwyer S, Rosman-Balzer DS, Swiss VA, et al. Treating Diet-Induced Diabetes and Obesity with Human Embryonic Stem Cell-Derived Pancreatic Progenitor Cells and Antidiabetic Drugs. Stem Cell Rep 2015; 4:605–20.

6. Rezania A, Bruin JE, Arora P, Rubin A, Batushansky I, Asadi A, O’Dwyer S, Quiskamp N, Mojibian M, Albrecht T, et al. Reversal of diabetes with insulin-producing cells derived in vitro from human pluripotent stem cells. Nat Biotechnol 2014; 32:1121–33.

7. Pagliuca FW, Millman JR, Gürtler M, Segel M, Van Dervort A, Ryu JH, Peterson QP, Greiner D, Melton DA. Generation of Functional Human Pancreatic β Cells In Vitro. Cell 2014; 159:428–39.

8. Kulkarni RN, Stewart AF. Summary of the Keystone Islet Workshop (April 2014): The Increasing Demand for Human Islet Availability in Diabetes Research. Diabetes 2014; 63:3979–81.

9. Manning Fox JE, Lyon J, Dai XQ, Wright RC, Hayward J, van de Bunt M, Kin T, Shapiro AMJ, McCarthy MI, Gloyn AL, et al. Human islet function following 20 years of cryogenic biobanking. Diabetologia 2015; 58:1503–12.

10. Nano R, Bosco D, Kerr-Conte JA, Karlsson M, Charvier S, Melzi R, Ezzouaoui R, Mercalli A, Hwa A, Pattou F, et al. Human islet distribution programme for basic research: activity over the last 5 years. Diabetologia 2015; 58:1138–40.

11. Kaddis JS, Olack BJ, Sowinski J, Cravens J, Contreras JL, Niland JC. Human pancreatic islets and diabetes research. JAMA 2009; 301:1580–7.

12. Lyon J, Manning Fox JE, Spigelman AF, Kim R, Smith N, O’Gorman D, Kin T, Shapiro AMJ, Rajotte RV, MacDonald PE. Research-Focused Isolation of Human Islets From Donors With and Without Diabetes at the Alberta Diabetes Institute IsletCore. Endocrinology 2016; 157:560–9.

13. ADI IsletCore. ADI IsletCore Homepage [accessed 2021 Nov 8]. http://www.bcell.org/adi-isletcore.html

14. ADI IsletCore. ADI IsletCore Webtool. [accessed 2021 Nov 8]. https://www.epicore.ualberta.ca/isletcore/Default

15. World Health Organization. Timeline: WHO’s COVID-19 response. [accessed 2021 Nov 8]. https://www.who.int/emergencies/diseases/novel-coronavirus-2019/interactive-timeline

16. Government of Alberta. Review of Alberta’s COVID-19 Pandemic Response: March 1 to October 12, 2020. https://www.alberta.ca/assets/documents/health-alberta-covid-19-pandemic-response-review-final-report.pdf

17. World Health Organization. WHO Coronavirus (COVID-19) Dashboard With Vaccination Data. [accessed 2021 Nov 8]. https://covid19.who.int/measures

18. Government of Alberta. COVID-19 Alberta statistics. [accessed 2021 Nov 8]. https://www.alberta.ca/covid-19-alberta-data.aspx

19. Ricordi C, Lacy PE, Finke EH, Olack B, Scharp D. Automated Method for Isolation of Human Pancreatic Islets. Diabetes 1988; 37:413–20.

20. Lyon J, Spigelman AF, Fox JEM, Macdonald PE. ADI IsletCore Protocols for the Isolation, Assessment and Cryopreservation of Human Pancreatic Islets of Langerhans for Research Purposes. protocols.io https://www.protocols.io/view/adi-isletcore-protocols-for-the-isolation-assessme-bupanvie

21. Wang L-J, Kin T, O’Gorman D, Shapiro AMJ, Naziruddin B, Takita M, Levy MF, Posselt AM, Szot GL, Savari O, et al. A Multicenter Study: North American Islet Donor Score in Donor Pancreas Selection for Human Islet Isolation for Transplantation. Cell Transplant 2016; 25:1515–23.

22. Kaddis JS, Danobeitia JS, Niland JC, Stiller T, Fernandez LA. Multi-Center Analysis of Novel and Established Variables Associated with Successful Human Islet Isolation Outcomes. Am J Transplant Off J Am Soc Transplant Am Soc Transpl Surg 2010; 10:646–56.

23. Zeng Y, Torre MA, Karrison T, Thistlethwaite JR. The correlation between donor characteristics and the success of human islet isolation. Transplantation 1994; 57:954–8.

24. O’Gorman D, Kin T, Murdoch T, Richer B, McGhee-Wilson D, Ryan EA, Shapiro JAM, Lakey JRT. The standardization of pancreatic donors for islet isolations. Transplantation 2005; 80:801–6.

25. Hilling DE, Bouwman E, Terpstra OT, Marang-van de Mheen PJ. Effects of donor-, pancreas-, and isolation-related variables on human islet isolation outcome: a systematic review. Cell Transplant 2014; 23:921–8.

26. Lakey JRT, Warnock GL, Rajotte RV, Suarez-Almazor ME, Ziliang AO, Shapiro AMJ, Kneteman NM. Variables in organ donors that affect the recovery of human islets of Langerhans. Transplantation 1996; 61:1047–53.

27. Ponte GM, Pileggi A, Messinger S, Alejandro A, Ichii H, Baidal DA, Khan A, Ricordi C, Goss JA, Alejandro R. Toward maximizing the success rates of human islet isolation: influence of donor and isolation factors. Cell Transplant 2007; 16:595–607.

28. Hanley SC, Paraskevas S, Rosenberg L. Donor and Isolation Variables Predicting Human Islet Isolation Success. Transplantation 2008; 85:950–5.

29. Ahmed O, Brockmeier D, Lee K, Chapman WC, Doyle MBM. Organ donation during the COVID-19 pandemic. Am J Transplant Off J Am Soc Transplant Am Soc Transpl Surg 2020; 20:3081–8.

30. Government of Canada. Travel Advice and Advisories [accessed 2021 Nov 9]. https://travel.gc.ca/travelling/advisories

31. Government of Canada. Aviation measures in response to COVID-19. [accessed 2021 Nov 9]. https://tc.canada.ca/en/initiatives/covid-19-measures-updates-guidance-issued-transport-canada/aviation-measures-response-covid-19

32. Andrew Freedman, John Muyskens, Chris Alcantara, Monica Ulmanu. How coronavirus grounded the airline industry. Wash. Post. 2020 Apr 01. [accessed 2021 Nov 9]. https://www.washingtonpost.com/graphics/2020/business/coronavirus-airline-industry-collapse/

33. Clayton HA, London NJ. Survival and function of islets during culture. Cell Transplant 1996; 5:1–12; discussion 13-17, 19.

34. Ichii H, Sakuma Y, Pileggi A, Fraker C, Alvarez A, Montelongo J, Szust J, Khan A, Inverardi L, Naziruddin B, et al. Shipment of Human Islets for Transplantation. Am J Transplant 2007; 7:1010–20.

35. Ikemoto T, Matsumoto S, Itoh T, Noguchi H, Tamura Y, Jackson AM, Shimoda M, Naziruddin B, Onaca N, Yasunami Y, et al. Assessment of islet quality following international shipping of more than 10,000 km. Cell Transplant 2010; 19:731–41.

36. Kaddis JS, Hanson MS, Cravens J, Qian D, Olack B, Antler M, Papas KK, Iglesias I, Barbaro B, Fernandez L, et al. Standardized transportation of human islets: an islet cell resource center study of more than 2,000 shipments. Cell Transplant 2013; 22:1101–11.

37. Poitout V, Satin LS, Kahn SE, Stoffers DA, Marchetti P, Gannon M, Verchere CB, Herold KC, Myers MG, Marshall SM. A Call for Improved Reporting of Human Islet Characteristics in Research Articles. Diabetes 2019; 68:239–40.

38. Poitout V, Satin LS, Kahn SE, Stoffers DA, Marchetti P, Gannon M, Verchere CB, Herold KC, Myers MG, Marshall SM. A call for improved reporting of human islet characteristics in research articles. Diabetologia 2019; 62:209–11.

39. Manning Fox JE, Spigelman AF, Dafoe T J. Alberta Diabetes Institute Welcome booklet: A user guide to human islets. [accessed 2021 Nov 21]. http://www.bcell.org/uploads/5/1/3/3/51338649/adi_isletcore_welcome_booklet.pdf

40. Friedman AL, Delli Carpini KW, Ezzell C, Irving H. There are no best practices in a pandemic: Organ donation within the COVID-19 epicenter. Am J Transplant Off J Am Soc Transplant Am Soc Transpl Surg 2020; 20:3089–93.

41. Azzi Y, Bartash R, Scalea J, Loarte-Campos P, Akalin E. COVID-19 and Solid Organ Transplantation: A Review Article. Transplantation 2021; 105:37–55.

42. Danziger-Isakov L, Blumberg EA, Manuel O, Sester M. Impact of COVID-19 in solid organ transplant recipients. Am J Transplant Off J Am Soc Transplant Am Soc Transpl Surg 2021; 21:925–37.

43. Olack BJ, Linetsky E, Niland JC. Human islet distribution during COVID-19 pandemic: the impact on diabetes research. CellR4 2020; 8: e2955

44. FedEx Canada. COVID-19 Service Updates. [accessed 2021 Nov 15]. https://www.fedex.com/en-ca/coronavirus.html

45. Santos T dos, Galipeau M, Gomes AS, Greenberg M, Larsen M, Lee D, Maghera J, Mulchandani CM, Patton M, Perera I, et al. Islet Biology during COVID-19: Progress and Perspectives. Can J Diabetes. Forthcoming.

